## Supplemental Table S2 for "Upscaling the surveillance of tick-borne pathogens in the French Caribbean islands"

**Table S2.** List of the positive control samples used for the Biomark system development.

| **Positive control** | **Sample type** | **Source** |
| --- | --- | --- |
| *Borrelia burgdorferi* | Culture (Strain B31) | National Reference Center of Borrelia (Strasbourg, France) |
| *Borrelia garinii* | Culture (Strain N11) | Lise Gern, University of Neuchâtel (Neuchâtel, Switzerland) |
| *Borrelia afzelii* | Culture (Strain NE632) | Lise Gern, University of Neuchâtel (Neuchâtel, Switzerland) |
| *Borrelia lusitaniae* | Culture (Strain Poti-B1) | Lise Gern, University of Neuchâtel (Neuchâtel, Switzerland) |
| *Borrelia lonestari* | Infected *Amblyomma americanum* (Tick collected in USA) | CDC (Atlanta, USA) |
| *Borrelia bissettii* | Plasmid, rpoB gene & pBluescriptIISK+ | GeneCust, September 2012 (Paris, France) |
| *Borrelia anserina* | Plasmid, fla gene & pBluescriptIISK+ | GeneCust, December 2015 (Paris, France) |
| *Borrelia parkeri* | Culture | Cecilia Hizo-Teufel, National Reference Center of Borrelia (Germany) |
| *Borrelia recurrentis* | Culture | ANSES (Maisons-Alfort, France) |
| *Anaplasma marginale* | Experimentally infected cow blood | Isabel Garcia Fernandez de Mera (Spain) |
| *Anaplasma phagocytophilum* | Infected *Ixodes* spp. (Tick collected in USA) | CDC (Atlanta, USA) |
| *Anaplasma platys* | Infected dog blood | H.J. Boulouis, ANSES (Maisons-Alfort, France) |
| *Anaplasma ovis* | Plasmid, msp4 gene & pBluescriptIISK+ | GeneCust, September 2012 (Paris, France) |
| *Anaplasma bovis* | Plasmid, groEL gene & pBluescriptIISK+ | GeneCust, January 2016 (Paris, France) |
| *Ehrlichia ewingii* | Infected *Amblyomma americanum* (Tick collected in USA) | CDC (Atlanta, USA) |
| *Ehrlichia chaffensis* | Infected *Amblyomma americanum* (Tick collected in USA) | CDC (Atlanta, USA) |
| *Ehrlichia ruminantium* | Culture | CIRAD (Guadeloupe, France) |
| *Panola mountain Ehrlichia* | Infected *Amblyomma americanum* (Tick collected in USA) | CDC (Atlanta, USA) |
| *Ehrlichia canis* | Culture | ANSES (Maisons-Alfort, France) |
| *Neoehrlichia mikurensis* | Infected rodent blood | Lars, Raberg, University of Lund (Sweden) |
| *Rickettsia conorii* | Infected *Rhipicephalus sanguineus* (Tick collected in USA) | CDC (Atlanta, USA) |
| *Rickettsia slovaca* | Culture | National Reference Center of Rickettsia (Marseille, France) |
| *Rickettsia massiliae* | Culture | National Reference Center of Rickettsia (Marseille, France) |
| *Rickettsia ricketsii* | Plasmid, ITS & pBluescriptIISK+ | GeneCust, Janvier 2016 (Paris, France) |
| *Rickettsia africae* | Culture | National Reference Center of Rickettsia (Marseille, France) |
| *Rickettsia typhi* | Culture | ANSES (Maisons-Alfort, France) |
| *Rickettsia prowazekii* | Plasmid, gltA gene & pBluescriptIISK+ | GeneCust, December 2016 (Paris, France) |
| *Rickettsia felis* | Culture | National Reference Center of Rickettsia (Marseille, France) |
| *Bartonella henselae* | Culture | ANSES (Maisons-Alfort, France) |
| *Bartonella quintana* | Culture | H.J. Boulouis, ANSES (Maisons-Alfort, France) |
| *Bartonella bacilliformis* | Culture | ANSES (Maisons-Alfort, France) |
| *Bartonella vinsonii subsp berkhoffii* | Culture | Lynn Osikowicz, stacey Bartlett, CDC (Colorado, USA) |
| *Francisella tularensis* | Culture | Nora Madani, ANSES (Maisons-Alfort, France) |
| *Coxiella burnetti* | Culture | Elodie Rousset, ANSES (Maisons-Alfort, France) |
| *Aegyptianella pullorum* | Plasmid, groEL gene & pBluescriptIISK+ | GeneCust, December 2015 (Paris, France) |
| *Babesia divergens* | Culture (clone RFS) | Laurence Malandrin, Oniris (Nantes, France) |
| *Babesia microti* | Culture (isolate R1) | Emmanuel Cornillot, University of Montpellier (Montpellier, France) |
| *Babesia bovis* | Plasmid, CCTeta gene & pBluescriptIISK+ | GeneCust, Janvier 2016 (Paris, France) |
| *Babesia canis* | Infected dog blood | VetAgroSup (Lyon, France) |
| *Babesia vogeli* | Infected dog blood | VetAgroSup (Lyon, France) |
| *Babesia Rossi* | Infected dog blood | ANSES (Maisons-Alfort, France) |
| *Babesia caballi* | Plasmid, Rap1 gene & pBluescriptIISK+ | GeneCust, September 2012 (Paris, France) |
| *Babesia bigemina* | Plasmid, 18S rRNA gene & pBluescriptIISK+ | GeneCust, September 2012 (Paris, France) |
| *Babesia ovis* | Plasmid, 18S rRNA gene & pBluescriptIISK+ | GeneCust, September 2012 (Paris, France) |
| *Babesia duncani* | Plasmid, ITS2 & pBluescriptIISK+ | GeneCust, January 2016 (Paris, France) |
| *Babesia gibsoni* | Plasmid, rap gene & pBluescriptIISK+ | GeneCust, December 2015 (Paris, France) |
| *Theileria parva* | Culture | Dirk Dobbelaera, University of Bern (Bern, Switzerland) |
| *Theileria annulata* | Culture | Dirk Dobbelaera, University of Bern (Bern, Switzerland) |
| *Theileria Lestoquari* | Culture | Dirk Dobbelaera, University of Bern (Bern, Switzerland) |
| *Cytauxzoon felis* | Plasmid, 18S rRNA gene & pBluescriptIISK+ | GeneCust, December 2015 (Paris, France) |
| *Hepatozoon canis* | Infected dog blood | Domenico Otranto, Alessio Giannelli, University of Bari (Bari, Italia) |
| *Leishmania infantum (chagasi)* | Culture | KASBARI Mohamed, ENVA (Maisons-Alfort, France) |
| *Leishmania Martiniquensis* | Culture | KASBARI Mohamed, ENVA (Maisons-Alfort, France) |
| *Rangelia vitalii* | Plasmid, 18S rRNA gene & pBluescriptIISK+ | GeneCust, December 2015 (Paris, France) |
| *Borrelia theileri* | Plasmid, glpQ gene & pBluescriptIISK+ | GeneCust, December 2017 (Paris, France) |
| *Theileria mutans* | Plasmid, ITS & pBluescriptIISK+ | GeneCust, December 2017 (Paris, France) |
| *Theileria velifera* | Plasmid, 18S rRNA gene & pBluescriptIISK+ | GeneCust, December 2017 (Paris, France) |
| *Theileria equi* | Plasmid, ema1 gene & pBluescriptIISK+ | GeneCust, December 2017 (Paris, France) |
| *Hepatozoon americanum* | Plasmid, 18S rRNA gene & pBluescriptIISK+ | GeneCust, December 2017 (Paris, France) |
| *Amblyomma variegatum* | Tick DNA (Tick collected in Guadeloupe) | Emmanuel Albina, CIRAD (Guadeloupe, France) |
| *Rhipicephalus microplus* | Tick DNA (Tick collected in Gualapagos) | ANSES (Maisons-Alfort, France) |
| *Rhipicephalus sanguineus* s.l. | Tick DNA (Tick collected in France) | ANSES (Maisons-Alfort, France) |
| *Borrelia burgdorferi* | Culture (Strain B31) | National Reference Center of Borrelia (Strasbourg, France) |
| *Borrelia garinii* | Culture (Strain N11) | Lise Gern, University of Neuchâtel (Neuchâtel, Switzerland) |
| *Borrelia afzelii* | Culture (Strain NE632) | Lise Gern, University of Neuchâtel (Neuchâtel, Switzerland) |
| *Borrelia lusitaniae* | Culture (Strain Poti-B1) | Lise Gern, University of Neuchâtel (Neuchâtel, Switzerland) |
| *Borrelia lonestari* | Infected *Amblyomma americanum* (Tick collected in USA) | CDC (Atlanta, USA) |
| *Borrelia bissettii* | Plasmid, rpoB gene & pBluescriptIISK+ | GeneCust, September 2012 (Paris, France) |
| *Borrelia anserina* | Plasmid, fla gene & pBluescriptIISK+ | GeneCust, December 2015 (Paris, France) |
| *Borrelia parkeri* | Culture | Cecilia Hizo-Teufel, National Reference Center of Borrelia (Germany) |
| *Borrelia recurrentis* | Culture | ANSES (Maisons-Alfort, France) |
| *Anaplasma marginale* | Experimentally infected cow blood | Isabel Garcia Fernandez de Mera (Spain) |
| *Anaplasma phagocytophilum* | Infected *Ixodes* spp. (Tick collected in USA) | CDC (Atlanta, USA) |
| *Anaplasma platys* | Infected dog blood | H.J. Boulouis, ANSES (Maisons-Alfort, France) |
| *Anaplasma ovis* | Plasmid, msp4 gene & pBluescriptIISK+ | GeneCust, September 2012 (Paris, France) |
| *Anaplasma bovis* | Plasmid, groEL gene & pBluescriptIISK+ | GeneCust, January 2016 (Paris, France) |
| *Ehrlichia ewingii* | Infected *Amblyomma americanum* (Tick collected in USA) | CDC (Atlanta, USA) |
| *Ehrlichia chaffensis* | Infected *Amblyomma americanum* (Tick collected in USA) | CDC (Atlanta, USA) |
| *Ehrlichia ruminantium* | Culture | CIRAD (Guadeloupe, France) |
| *Panola mountain Ehrlichia* | Infected *Amblyomma americanum* (Tick collected in USA) | CDC (Atlanta, USA) |
| *Ehrlichia canis* | Culture | ANSES (Maisons-Alfort, France) |
| *Neoehrlichia mikurensis* | Infected rodent blood | Lars, Raberg, University of Lund (Sweden) |
| *Rickettsia conorii* | Infected *Rhipicephalus sanguineus* (Tick collected in USA) | CDC (Atlanta, USA) |
| *Rickettsia slovaca* | Culture | National Reference Center of Rickettsia (Marseille, France) |
| *Rickettsia massiliae* | Culture | National Reference Center of Rickettsia (Marseille, France) |
| *Rickettsia ricketsii* | Plasmid, ITS & pBluescriptIISK+ | GeneCust, Janvier 2016 (Paris, France) |
| *Rickettsia africae* | Culture | National Reference Center of Rickettsia (Marseille, France) |
| *Rickettsia typhi* | Culture | ANSES (Maisons-Alfort, France) |
| *Rickettsia prowazekii* | Plasmid, gltA gene & pBluescriptIISK+ | GeneCust, December 2016 (Paris, France) |
| *Rickettsia felis* | Culture | National Reference Center of Rickettsia (Marseille, France) |
| *Bartonella henselae* | Culture | ANSES (Maisons-Alfort, France) |
| *Bartonella quintana* | Culture | H.J. Boulouis, ANSES (Maisons-Alfort, France) |
| *Bartonella bacilliformis* | Culture | ANSES (Maisons-Alfort, France) |
| *Bartonella vinsonii subsp berkhoffii* | Culture | Lynn Osikowicz, stacey Bartlett, CDC (Colorado, USA) |
| *Francisella tularensis* | Culture | Nora Madani, ANSES (Maisons-Alfort, France) |
| *Coxiella burnetti* | Culture | Elodie Rousset, ANSES (Maisons-Alfort, France) |
| *Aegyptianella pullorum* | Plasmid, groEL gene & pBluescriptIISK+ | GeneCust, December 2015 (Paris, France) |
| *Babesia divergens* | Culture (clone RFS) | Laurence Malandrin, Oniris (Nantes, France) |
| *Babesia microti* | Culture (isolate R1) | Emmanuel Cornillot, University of Montpellier (Montpellier, France) |
| *Babesia bovis* | Plasmid, CCTeta gene & pBluescriptIISK+ | GeneCust, Janvier 2016 (Paris, France) |
| *Babesia canis* | Infected dog blood | VetAgroSup (Lyon, France) |
| *Babesia vogeli* | Infected dog blood | VetAgroSup (Lyon, France) |
| *Babesia Rossi* | Infected dog blood | ANSES (Maisons-Alfort, France) |
| *Babesia caballi* | Plasmid, Rap1 gene & pBluescriptIISK+ | GeneCust, September 2012 (Paris, France) |
| *Babesia bigemina* | Plasmid, 18S rRNA gene & pBluescriptIISK+ | GeneCust, September 2012 (Paris, France) |
| *Babesia ovis* | Plasmid, 18S rRNA gene & pBluescriptIISK+ | GeneCust, September 2012 (Paris, France) |
| *Babesia duncani* | Plasmid, ITS2 & pBluescriptIISK+ | GeneCust, January 2016 (Paris, France) |
| *Babesia gibsoni* | Plasmid, rap gene & pBluescriptIISK+ | GeneCust, December 2015 (Paris, France) |
| *Theileria parva* | Culture | Dirk Dobbelaera, University of Bern (Bern, Switzerland) |
| *Theileria annulata* | Culture | Dirk Dobbelaera, University of Bern (Bern, Switzerland) |
| *Theileria Lestoquari* | Culture | Dirk Dobbelaera, University of Bern (Bern, Switzerland) |
| *Cytauxzoon felis* | Plasmid, 18S rRNA gene & pBluescriptIISK+ | GeneCust, December 2015 (Paris, France) |
| *Hepatozoon canis* | Infected dog blood | Domenico Otranto, Alessio Giannelli, University of Bari (Bari, Italia) |
| *Leishmania infantum (chagasi)* | Culture | KASBARI Mohamed, ENVA (Maisons-Alfort, France) |
| *Leishmania Martiniquensis* | Culture | KASBARI Mohamed, ENVA (Maisons-Alfort, France) |
| *Rangelia vitalii* | Plasmid, 18S rRNA gene & pBluescriptIISK+ | GeneCust, December 2015 (Paris, France) |
| *Borrelia theileri* | Plasmid, glpQ gene & pBluescriptIISK+ | GeneCust, December 2017 (Paris, France) |
| *Theileria mutans* | Plasmid, ITS & pBluescriptIISK+ | GeneCust, December 2017 (Paris, France) |
| *Theileria velifera* | Plasmid, 18S rRNA gene & pBluescriptIISK+ | GeneCust, December 2017 (Paris, France) |
| *Theileria equi* | Plasmid, ema1 gene & pBluescriptIISK+ | GeneCust, December 2017 (Paris, France) |
| *Hepatozoon americanum* | Plasmid, 18S rRNA gene & pBluescriptIISK+ | GeneCust, December 2017 (Paris, France) |
| *Amblyomma variegatum* | Tick DNA (Tick collected in Guadeloupe) | Emmanuel Albina, CIRAD (Guadeloupe, France) |
| *Rhipicephalus microplus* | Tick DNA (Tick collected in Gualapagos) | ANSES (Maisons-Alfort, France) |
| *Rhipicephalus sanguineus* s.l. | Tick DNA (Tick collected in France) | ANSES (Maisons-Alfort, France) |
