## Supplemental Figure S1 for "Upscaling the surveillance of tick-borne pathogens in the French Caribbean islands"

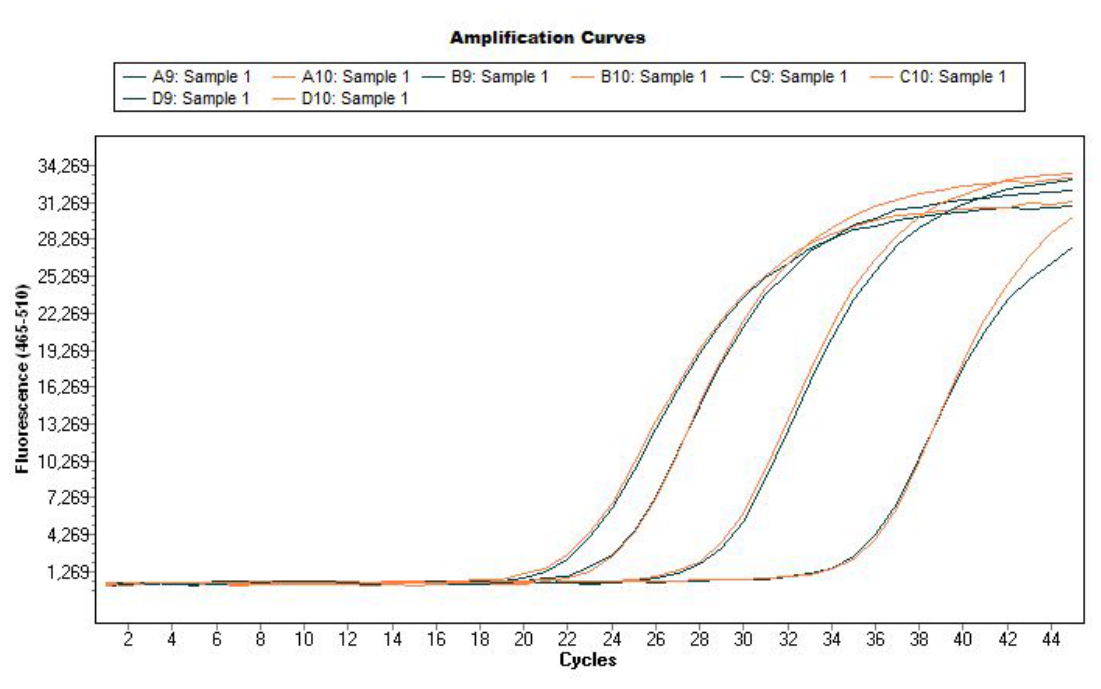

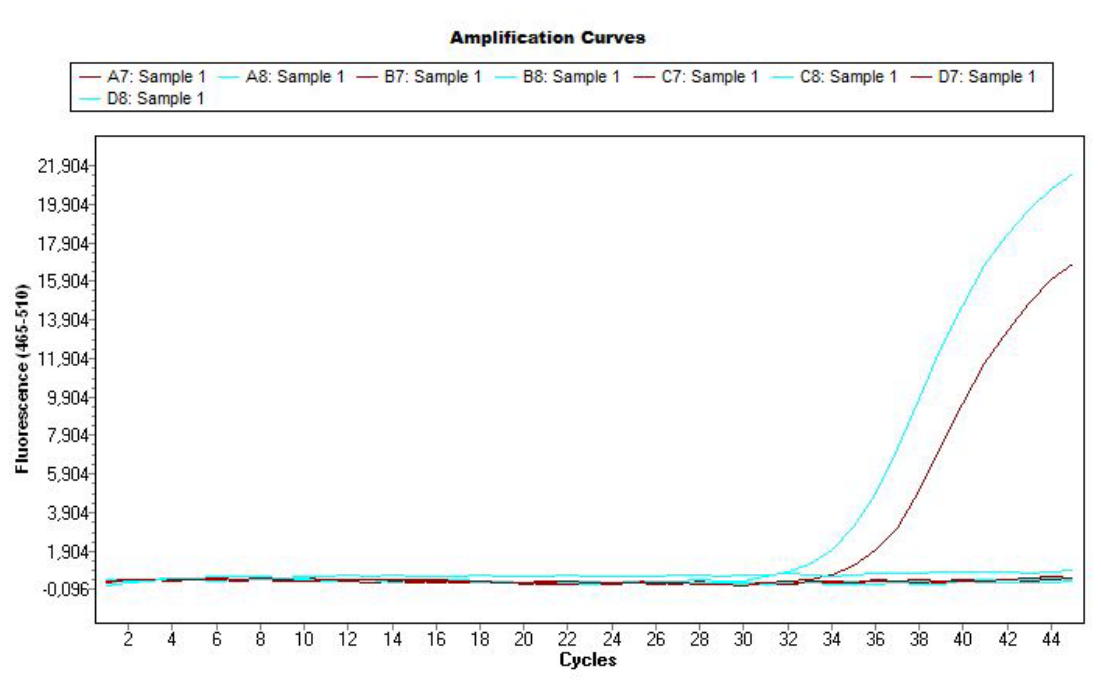

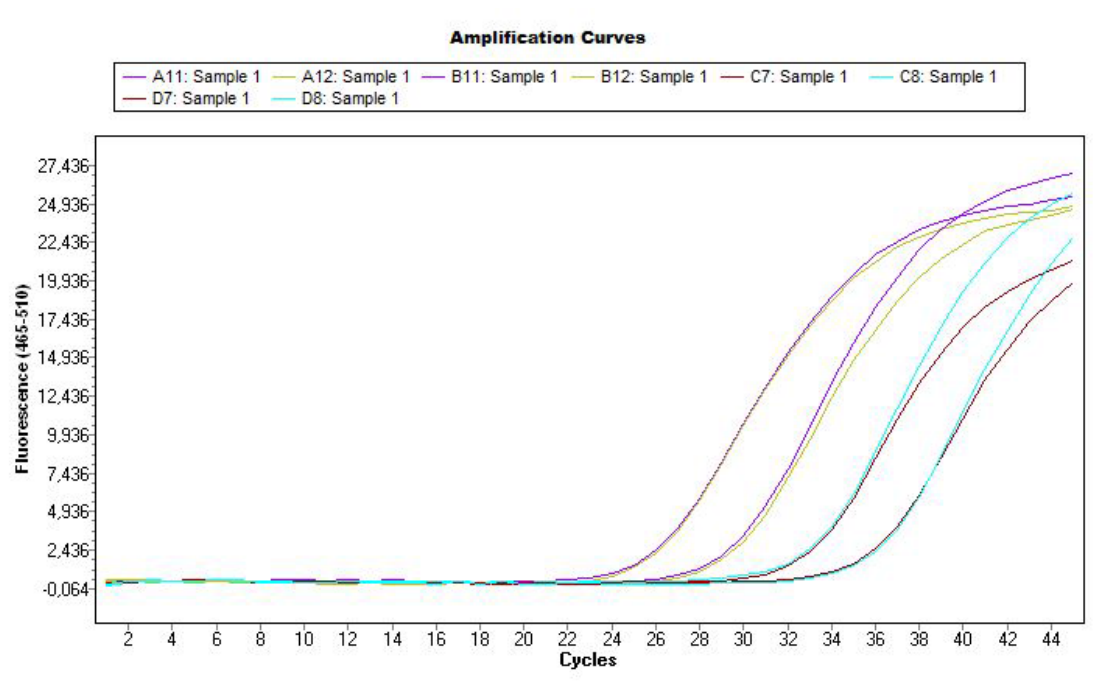

**(a)**

**(b)**

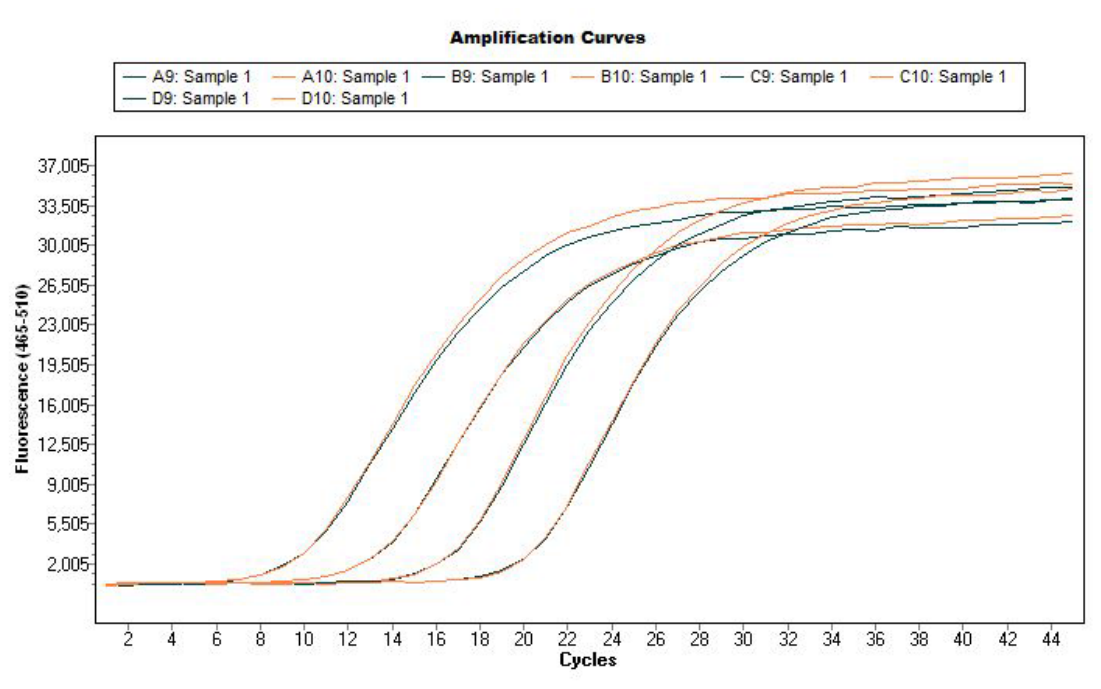

**(c)**

**(d)**

**Figure S1.** Improvement of detection signals by pre-amplification. Test of primer/probe set sensitivity for a range of dilutions of positive controls by TaqMan real-time PCR using LightCycler 480, before and after pre-amplification. Results of the sensitivity test of the *Leishmania infantum* design using *a Leishmania infantum* culture, before (a) and after (c) pre-amplification; Results of the sensitivity test of the *Rickettsia* spp. design using *Rickettsia conorii*-positive controls (extracted from an infected *Rhipicephalus sanguineus* tick), before (b) and after (d) pre-amplification.
