## Supplemental Table S1 for "Upscaling the surveillance of tick-borne pathogens in the French Caribbean islands"

**Table S1.** GPS coordinates of the tick collection sites and number of ticks collected. A total of 578 adult ticks collected from cattle from Guadeloupe and Martinique were used for the screening of tick-borne pathogens with the newly implemented BioMark™ real-time PCR system.

| **Location** | **Collection Sites** | **Cattle** | **Animal** | **Tick species** | **Tick number** |
| --- | --- | --- | --- | --- | --- |
| GUADELOUPE |  |  |  |  | 297 |
|  |  |  |  | *Amblyomma variegatum* | 132 |
|  |  |  |  | *Rhipicephalus microplus* | 165 |
|  | Sainte-Anne (N 16° 14'; W 61° 23') |  |  |  | 42 |
|  |  | Cattle 1 | Animal 1 | *Amblyomma variegatum* | 19 |
|  |  | Cattle 2 | Animal 2 | *Amblyomma variegatum* | 2 |
|  |  |  |  | *Rhipicephalus microplus* | 4 |
|  |  |  | Animal 3 | *Amblyomma variegatum* | 7 |
|  |  |  |  | *Rhipicephalus microplus* | 9 |
|  |  | Cattle 3 | Animal 4 | *Amblyomma variegatum* | 1 |
|  | Petit Bourg (N 16° 10'; W 61° 36') |  |  |  | 48 |
|  |  | Cattle 4 | Animal 5 | *Amblyomma variegatum* | 11 |
|  |  |  | Animal 6 | *Amblyomma variegatum* | 9 |
|  |  |  |  | *Rhipicephalus microplus* | 5 |
|  |  | Cattle 5 | Animal 7 | *Amblyomma variegatum* | 4 |
|  |  |  |  | *Rhipicephalus microplus* | 1 |
|  |  |  | Animal 8 | *Amblyomma variegatum* | 13 |
|  |  | Cattle 6 | Animal 9 | *Amblyomma variegatum* | 1 |
|  |  |  | Animal 10 | *Amblyomma variegatum* | 4 |
|  | Capesterre (N 16° 2'; W 61° 34') |  |  |  | 50 |
|  |  | Cattle 7 | Animal 11 | *Amblyomma variegatum* | 9 |
|  |  |  | Animal 12 | *Amblyomma variegatum* | 1 |
|  |  |  |  | *Rhipicephalus microplus* | 2 |
|  |  | Cattle 8 | Animal 13 | *Amblyomma variegatum* | 1 |
|  |  |  |  | *Rhipicephalus microplus* | 16 |
|  |  |  | Animal 14 | *Amblyomma variegatum* | 6 |
|  |  |  |  | *Rhipicephalus microplus* | 1 |
|  |  | Cattle 9 | Animal 15 | *Amblyomma variegatum* | 2 |
|  |  |  |  | *Rhipicephalus microplus* | 5 |
|  |  |  | Animal 16 | *Rhipicephalus microplus* | 7 |
|  | Le Gosier (N 16° 13'; W 61° 28') |  |  |  | 27 |
|  |  | Cattle 10 | Animal 17 | *Amblyomma variegatum* | 11 |
|  |  |  |  | *Rhipicephalus microplus* | 3 |
|  |  |  | Animal 18 | *Amblyomma variegatum* | 7 |
|  |  | Cattle 11 | Animal 19 | *Amblyomma variegatum* | 1 |
|  |  |  | Animal 20 | *Amblyomma variegatum* | 3 |
|  |  | Cattle 12 | Animal 21 | *Amblyomma variegatum* | 2 |
|  | Pointe-Noire (N 16° 13'; W 61° 45') |  |  |  | 13 |
|  |  | Cattle 13 | Animal 22 | *Amblyomma variegatum* | 2 |
|  |  |  | Animal 23 | *Amblyomma variegatum* | 2 |
|  |  |  | Animal 24 | *Amblyomma variegatum* | 4 |
|  |  |  | Animal 25 | *Amblyomma variegatum* | 2 |
|  |  | Cattle 14 | Animal 26 | *Amblyomma variegatum* | 2 |
|  |  |  | Animal 27 | *Amblyomma variegatum* | 1 |
|  | Les Abymes (N 16° 16'; W 61° 31') |  |  |  | 68 |
|  |  | Cattle 15 | Animal 28 | *Amblyomma variegatum* | 1 |
|  |  | Cattle 16 | Animal 29 | *Rhipicephalus microplus* | 10 |
|  |  |  | Animal 30 | *Rhipicephalus microplus* | 6 |
|  |  | Cattle 17 | Animal 31 | *Rhipicephalus microplus* | 18 |
|  |  |  | Animal 32 | *Rhipicephalus microplus* | 31 |
|  |  | Cattle 18 | Animal 33 | *Rhipicephalus microplus* | 2 |
|  | Morne-À-l'Eau (N 16° 19'; W 61° 28') |  |  |  | 46 |
|  |  | Cattle 19 | Animal 34 | *Rhipicephalus microplus* | 6 |
|  |  |  | Animal 35 | *Rhipicephalus microplus* | 2 |
|  |  | Cattle 20 | Animal 36 | *Rhipicephalus microplus* | 19 |
|  |  |  | Animal 37 | *Rhipicephalus microplus* | 7 |
|  |  | Cattle 21 | Animal 38 | *Amblyomma variegatum* | 1 |
|  |  |  |  | *Rhipicephalus microplus* | 11 |
|  | Bouillante (N 16° 7'; W 61° 46') |  |  |  | 3 |
|  |  | Cattle 22 | Animal 39 | *Amblyomma variegatum* | 2 |
|  |  |  | Animal 40 | *Amblyomma variegatum* | 1 |
| MARTINIQUE |  |  |  |  | 281 |
|  |  |  |  | *Rhipicephalus microplus* | 281 |
|  | Rivière-Pilote (N 14° 28', W 60° 54') |  |  |  | 4 |
|  |  | Cattle 1 | Animal 1 | *Rhipicephalus microplus* | 4 |
|  | Le Diamant (N 14° 29'; W 61° 1') |  |  |  | 1 |
|  |  | Cattle 2 | Animal 2 | *Rhipicephalus microplus* | 1 |
|  | Le François (N 14° 36'; W 60° 53') |  |  |  | 52 |
|  |  | Cattle 3 | Animal 3 | *Rhipicephalus microplus* | 50 |
|  |  | Cattle 4 | Animal 4 | *Rhipicephalus microplus* | 2 |
|  | Le Lamentin (N 14° 37'; W 60° 59') |  |  |  | 2 |
|  |  | Cattle 5 | Animal 5 | *Rhipicephalus microplus* | 2 |
|  | Le Lorrain (N 14° 49'; W 61° 2') |  |  |  | 5 |
|  |  | Cattle 6 | Animal 6 | *Rhipicephalus microplus* | 5 |
|  | Le Morne-Vert (N 14° 42'; W 61° 8') |  |  |  | 3 |
|  |  | Cattle 7 | Animal 7 | *Rhipicephalus microplus* | 3 |
|  | Le Robert (N 14° 40'; W 60° 56') |  |  |  | 57 |
|  |  | Cattle 8 | Animal 8 | *Rhipicephalus microplus* | 47 |
|  |  | Cattle 9 | Animal 9 | *Rhipicephalus microplus* | 3 |
|  |  | Cattle 10 | Animal 10 | *Rhipicephalus microplus* | 6 |
|  |  | Cattle 11 | Animal 11 | *Rhipicephalus microplus* | 1 |
|  | Le Vauclin (N 14° 32'; W 60° 50') |  |  |  | 40 |
|  |  | Cattle 12 | Animal 12 | *Rhipicephalus microplus* | 12 |
|  |  | Cattle 13 | Animal 13 | *Rhipicephalus microplus* | 6 |
|  |  | Cattle 14 | Animal 14 | *Rhipicephalus microplus* | 3 |
|  |  | Cattle 15 | Animal 15 | *Rhipicephalus microplus* | 4 |
|  |  | Cattle 16 | Animal 16 | *Rhipicephalus microplus* | 2 |
|  |  | Cattle 17 | Animal 17 | *Rhipicephalus microplus* | 5 |
|  |  | Cattle 18 | Animal 18 | *Rhipicephalus microplus* | 3 |
|  |  | Cattle 19 | Animal 19 | *Rhipicephalus microplus* | 5 |
|  | Rivière-Salée (N 14° 31'; W 60° 57') |  |  |  | 3 |
|  |  | Cattle 20 | Animal 20 | *Rhipicephalus microplus* | 3 |
|  | Saint-Esprit (N 14° 33'; W 60° 55') |  |  |  | 28 |
|  |  | Cattle 21 | Animal 21 | *Rhipicephalus microplus* | 28 |
|  | Saint-Joseph (N 14° 40'; W 61° 2') |  |  |  | 13 |
|  |  | Cattle 22 | Animal 22 | *Rhipicephalus microplus* | 7 |
|  |  | Cattle 23 | Animal 23 | *Rhipicephalus microplus* | 6 |
|  | Sainte-Anne (N 14° 26'; W 60° 50') |  |  |  | 29 |
|  |  | Cattle 24 | Animal 24 | *Rhipicephalus microplus* | 3 |
|  |  | Cattle 25 | Animal 25 | *Rhipicephalus microplus* | 8 |
|  |  | Cattle 26 | Animal 26 | *Rhipicephalus microplus* | 18 |
|  | Sainte-Luce (N 14° 29'; W 60° 56') |  |  |  | 36 |
|  |  | Cattle 27 | Animal 27 | *Rhipicephalus microplus* | 3 |
|  |  | Cattle 28 | Animal 28 | *Rhipicephalus microplus* | 33 |
|  | Sainte-Marie (N 14° 46'; W 60° 59') |  |  |  | 8 |
|  |  | Cattle 29 | Animal 29 | *Rhipicephalus microplus* | 8 |
| Total |  |  |  |  | 578 |
